# Mapping molecular diffusion across the whole cell with spatial statistics-based FRAP

**DOI:** 10.1101/2025.02.08.637213

**Authors:** Yohei Okabe, Takumi Saito, Outa Nakashima, Daiki Matsunaga, Shinji Deguchi

## Abstract

Understanding molecular diffusion within cells is crucial for gaining insights into cellular biophysical mechanisms. Fluorescence recovery after photobleaching (FRAP) is a powerful technique for assessing molecular diffusion, yet its localized measurement approach hinders whole-cell analysis. To overcome this limitation, we developed Probabilistic FRAP (Pro-FRAP), a novel approach that integrates FRAP with sequential Gaussian simulation (SGS), an advanced spatial statistical method incorporating probabilistic modeling to estimate diffusion in unmeasured regions. Pro-FRAP applies SGS to standardize measured FRAP data, perform conditional simulations based on spatial correlations, and generate statistically robust estimates. In separate analyses, numerical simulations were conducted to optimize the spatial arrangement of measurement points, enhancing data accuracy and coverage. Unlike deterministic interpolation methods, Pro-FRAP captures spatial variability and quantifies uncertainty in intracellular diffusion, providing a more detailed representation of molecular transport. Thus, this framework extends FRAP beyond localized measurements, offering a refined approach for mapping intracellular diffusion with improved spatial coverage and statistical reliability.

**Significance statement:** Understanding molecular diffusion is essential for deciphering cellular functions and regulation. However, conventional fluorescence recovery after photobleaching (FRAP) techniques are constrained to localized measurements, limiting their ability to capture diffusion dynamics across entire cells. Here, we describe Probabilistic FRAP (Pro-FRAP), a novel framework that integrates FRAP with sequential Gaussian simulation (SGS), a spatial statistical method that enables the estimation of molecular diffusion in unmeasured regions. By incorporating probabilistic modeling, Pro-FRAP generates high-resolution diffusion maps with enhanced statistical reliability. This approach provides a more comprehensive perspective on intracellular transport, allowing for deeper insights into the spatial organization of cellular processes.

## Introduction

Understanding molecular diffusion within cells is crucial for gaining insight into the complex biophysical mechanisms that maintain cellular homeostasis. Molecular diffusion facilitates the transport of signaling molecules, nutrients, and other essential compounds, ensuring cellular functionality and adaptability. Probing diffusion rates and patterns advances our comprehension of cellular physiology, stress responses, disease mechanisms, and therapeutic effects.

Fluorescence recovery after photobleaching (FRAP) is a powerful technique for evaluating molecular diffusion in living cells (1–5). In FRAP measurements, defined regions of a cell are photobleached, and the subsequent recovery of fluorescence is monitored over time. This recovery indicates the movement of fluorescently labeled molecules within the bleached region, allowing for the quantification of diffusion coefficients. FRAP has been pivotal in revealing intracellular dynamics, providing quantitative insights into the local mobility and functional roles of molecules. Its broad applicability across different cell types and organisms (6, 7) has made FRAP an indispensable tool in biology research.

Despite its significant contributions, FRAP is inherently limited by its localized measurement approach, which poses challenges for whole-cell analysis. Specifically, conducting FRAP measurements at multiple sites within a single cell can lead to excessive photobleaching, reducing overall fluorescence intensity and therefore compromising measurement accuracy. This practical constraint confines FRAP to discrete regions, limiting the acquisition of comprehensive spatial data across the entire cell. Consequently, capturing the full complexity of molecular dynamics within the cellular context becomes difficult, preventing a thorough analysis. This limitation is particularly significant when studying processes that involve extensive spatial variation, such as intracellular transport and membrane trafficking, where understanding the global behavior is crucial.

To address these limitations, we propose Probabilistic FRAP (Pro-FRAP), a novel approach that integrates FRAP with sequential Gaussian simulation (SGS), an advanced spatial statistical method, to estimate molecular diffusion across unmeasured regions. Unlike deterministic methods such as interpolation through averaging between measurement points (8, 9), SGS is a probabilistic simulation technique that generates multiple realizations incorporating spatial variability, enabling an uncertainty-aware assessment of diffusion patterns (10–12). By integrating the spatial statistics with FRAP, Pro-FRAP extends conventional diffusion analysis beyond localized measurements, providing statistically robust and spatially continuous estimates of the intracellular diffusion landscape.

## 2. Material and methods

### 2.1 Cell culture

U2OS cells (HTB-96, ATCC) were cultured in high-glucose DMEM (Wako) supplemented with 10% fetal bovine serum (Sigma), 1% penicillin-streptomycin (Wako), and 200 mM L-Glutamine, and maintained at 37°C in a humidified 5% CO_2_ incubator. Fluorescent probes, 5-chloromethylfluorescein diacetate (CellTracker-Green, Invitrogen), were diluted to 1 mM with DMSO to prepare a stock solution. Cells were incubated in Opti-MEM containing 5-chloromethylfluorescein diacetate at a final concentration of 5 μM for 30 min at 37°C. For experiments, cells were incubated in FluoroBright (Gibco) supplemented with 1M HEPES, 10% fetal bovine serum (Sigma), 1% penicillin-streptomycin (Wako), 200 mM L-glutamine, and 100 mM sodium pyruvate.

### 2.2 FRAP experiments

FRAP experiments were performed with a confocal laser scanning microscope (FV3000, Olympus) mounting a 60x/NA1.42 objective. To increase the frame rate, the entire image of a single cell was divided into four segments, which were sequentially captured. Within each segment, images were acquired using a 488 nm excitation laser, and multiple circular regions spaced ~10 μm apart were photobleached using 405- and 488-nm lasers. The temporal evolution of fluorescence intensity distribution within each circular region *F*(*r, t*) was modeled with a Gaussian function

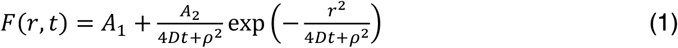

where *D* is the diffusion coefficient, and *A*_1_, *A*_2_, and *ρ* are fitting parameters, all determined using the least-square method (6, 13, 14).

### 2.3 Ordinary kriging

To estimate spatially distributed *D* values from measured data points, ordinary kriging was employed as a spatial statistics-based interpolation method (15). This approach provides optimal linear unbiased predictions by incorporating the spatial autocorrelation of the data to estimate unknown values with minimal error. Specifically, ordinary kriging utilizes a semi-variogram to quantify how spatial variance varies with the distance between data points, enabling statistically robust interpolation even in heterogeneous datasets. The predicted value at an unobserved location *o* is expressed as a linear combination of values observed at locations *k*,

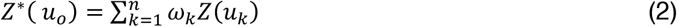

where the kriging weights *ω*_*k*_ represent the contributions of each observed value *Z* at position *u*_*k*_; *u*_*o*_ denotes the position of an unobserved point; an asterisk indicates estimated variables; and *n* is the total number of observed data points. The weights *ω*_*k*_ are determined by solving the following kriging equations,

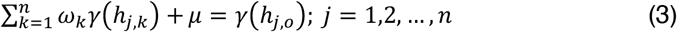

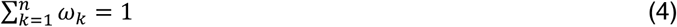

where the terms *γ*(*h*_*j,k*_) and *γ*(*h*_*j,o*_) are the semi-variogram for the distance between observation positions *u*_*j*_ and *u*_*k*_ and between observation positions *u*_*j*_ and the prediction position *u*_*o*_, respectively, and the parameter *µ* is the Lagrange multiplier ensuring unbiased predictions (see Supporting note 1 for details). The semi-variogram itself is calculated directly from the raw data of the FRAP experiments *Z*,

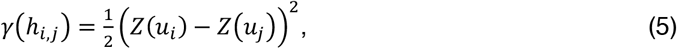

and is characterized by fitting a widely used exponential model to the variogram cloud, consisting of all data,

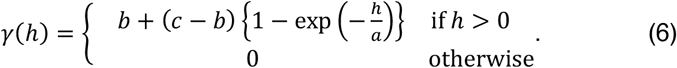

By minimizing the prediction variance 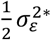, where the factor 1/2 is included to simplify quadratic terms, ordinary kriging ensures the best linear unbiased prediction while considering spatial autocorrelation and measurement uncertainty. Solving the resulting kriging equations provides both the predicted values and the associated prediction uncertainty.

### 2.4 Spatial autocorrelation analysis

Moran’s *I*, defined as

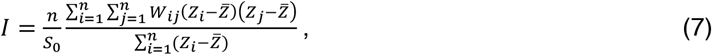

was employed to evaluate the spatial autocorrelation of dataset, ensuring the robustness of the interpolation results, in which 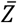 is the mean of the data, and *W*_*ij*_ represents the spatial weight between data points *i* and *j. W*_*ij*_ and *S*_0_ are defined as

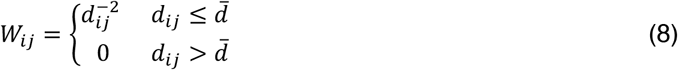

and

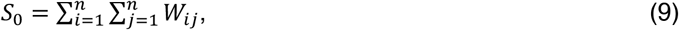

respectively, where *d*_*ij*_ is the distance between *u*_*i*_ and *u*_*j*_, and 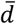 is the mean of *d*_*ij*_. To statistically test the spatial correlation, the *z*-score is defined as

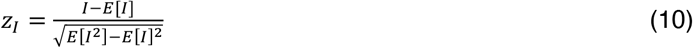

where *E* represents the expected value. The null hypothesis assuming no spatial autocorrelation is tested using the *p*-value derived from the *z*-score and the standard normal distribution. A *p*-value less than 0.01 was considered statistically significant, indicating the presence of spatial autocorrelation in the dataset.

### 2.5 Sequential Gaussian simulation

Pro-FRAP utilizes sequential Gaussian simulation (SGS), a probabilistic statistical method used to generate spatially distributed values that preserve spatial variability while closely matching observed data. While kriging provides a single optimal estimate based on spatial interpolation, SGS employs kriging as part of a stochastic simulation process, generating multiple realizations to represent plausible spatial distributions and quantify uncertainty. To ensure that the simulation assumptions hold, the cumulative probability distribution of the diffusion coefficients obtained from FRAP experiments was first examined. A normal score transformation was applied to all datasets to standardize the data. To reduce computational load, the original data captured at a resolution of 300 × 300 was downsampled to 70 × 70 using a moving average before computation. A grid for estimation points was then constructed based on the observed cell morphology, restricting the estimation to intracellular regions. The sequence of estimation points, which strongly influences the simulation outcome, was determined randomly using uniform random numbers to minimize systematic biases. At each estimation point, ordinary kriging was used to compute the best linear unbiased estimator along with its corresponding error variance. A random value drawn from a normal distribution with variance equal to the kriging error variance was then added to account for spatial uncertainty. These estimated values were sequentially incorporated into the dataset, ensuring that each subsequent estimation was conditioned on the updated dataset, thus preserving spatial consistency. For datasets that had undergone normal score transformation, an inverse transformation was applied at the final stage to return the estimated values to their original scale. The simulation process was repeated 100 times within the Pro-FRAP framework, generating multiple realizations for each estimation point. The mean of these realizations was taken as the final estimated value, while the spread of these realizations provided a measure of spatial variability and confidence in the predictions.

### 2.6 Analysis of intracellular diffusion coefficients

The intracellular diffusion coefficients were analyzed based on the fluorescence intensity of 5-chloromethylfluorescein diacetate, which reflects the degree of probe uptake. To achieve this, thresholding was applied using Fiji/ImageJ software (National Institutes of Health) to automatically segment and extract regions. Intracellular regions were classified into the nucleoplasm, endoplasmic reticulum (ER), and the rest of the cytoplasm. Data were collected from *N* = 4 independent experiments, and statistical comparisons of diffusion coefficients across categories were performed. Statistical significance was assessed using an unpaired two-tailed Student’s *t*-test for raw experimental data before interpolation and using a one-way analysis of variance for data after interpolation. Statistical significance was defined as follows: ***, *p* ≤ 0.001.

### 2.7 Simulated evaluation of spatial sampling strategies

To identify optimal spatial sampling patterns for accurately estimating intracellular diffusion coefficients, we simulated interpolation processes across multiple spatial configurations. The analysis considered three key factors: (i) the spatial arrangement of measurement points, (ii) the underlying diffusion coefficient distribution, and (iii) cellular morphology. First, two spatial sampling patterns were examined: a Gaussian-distributed pattern, where photobleached positions were assigned based on a Gaussian function with varying standard deviations *σ* centered within the cell, and a lattice-based pattern, where measurement points were arranged in a grid-like structure. Second, diffusion coefficient distributions were modeled in two ways: a radial gradient with concentric contours, where diffusion coefficients gradually changed with distance from the center, and a stepwise variation, where diffusion coefficients exhibited discrete jumps between concentric regions rather than a smooth gradient. Third, two distinct cellular morphologies, derived from experimental images, were analyzed to assess the impact of cell shape on prediction accuracy. To evaluate the influence of these factors, interpolated diffusion maps generated by Pro-FRAP were compared against the original ground truth using two error metrics: mean absolute error (MAE) and root mean squared error (RMSE).

## 3. Results

### 3.1 Whole-cell diffusion mapping with Pro-FRAP

We performed FRAP on the fluorescent probe, 5-chloromethylfluorescein diacetate, in U2OS cells (Fig. 1A). Photobleaching was performed at multiple sites approximately equidistantly distributed throughout individual cells. To determine the diffusion coefficients in each photobleached region, spatial and temporal intensity changes were analyzed using the least-square method with a Gaussian model (Fig. 1B-1D). The diffusion coefficients were then obtained as a discrete set of values (Fig. 1E) and subsequently utilized as a dataset to calculate Moran’s *I*, a measure of spatial correlation. Using a spatial weights matrix to define relationships between data points based on their spatial proximity, the associated *p*-value was determined by comparing the observed Moran’s *I* to its expected value under the null hypothesis of no spatial correlation (Fig. 1B). A semi-variogram was constructed to represent variance as a function of distance, and a widely used exponential model was fitted to determine the spatial structure parameters (Fig. 1F) (15). The parameter *a* in Eq. (6), referred to as the “range” in the exponential model and representing the distance within which spatial correlation is significant, was found in many cases to exceed the characteristic length scale of individual cells, i.e. ~30–50 μm, indicating that spatial correlation was present within the cell. Subsequently, diffusion coefficients at each pixel within cells were determined through the process of SGS (Fig. 1G). Many of these datasets exhibited spatial correlation, as indicated by high Moran’s *I* values (>0.3) and low *p*-values (<0.001) (Fig. S1).

**Figure 1.**
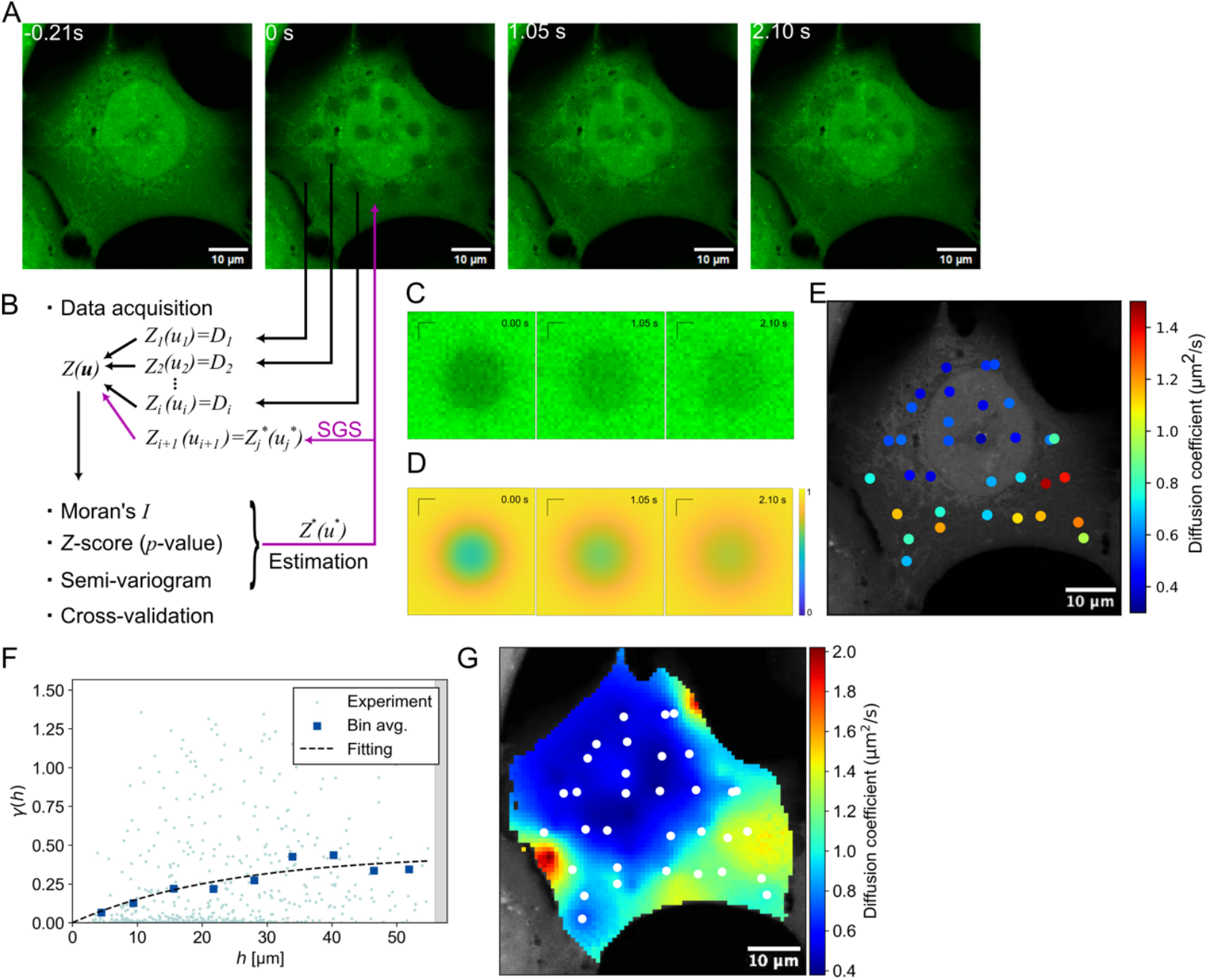
Demonstration of Pro-FRAP. (A) FRAP experiments on fluorescent probes, 5-chloromethylfluorescein diacetate in a U2OS cell. (B) Schematics of workflow in spatial statics. (C, D) Time course of a representative FRAP region in experiments (C) and regression analysis (D). (E) Diffusion coefficients measured at each FRAP region. (F) The semi-variogram of each data pair (blue dots), data binning (blue rectangle), and a regression curve (dashed line). (G) Diffusion coefficients determined by Pro-FRAP. White dots represent measurement points. Scale: 1 μm in C, D; 10 μm in others.

To evaluate the effectiveness of our approach, we conducted a cross-validation analysis (Fig. S1). Cross-validation is an essential step in assessing the validity and reliability of predictions. By dividing the data into subsets for training and testing, this method ensures that predictions are evaluated using unseen data, providing an unbiased estimate of model performance and minimizing the risk of overfitting. The analysis showed that the SGS-based approach consistently yielded more accurate estimates than ordinary kriging alone, a widely used method in conventional spatial statistics, across all four representative datasets. Specifically, the prediction accuracies for ordinary kriging alone were 71%, 68%, 85%, and 78%, while the incorporation of SGS improved these values to 75%, 82%, 91%, and 93%, respectively, demonstrating that SGS enhances predictive performance by effectively capturing spatial variability. In separate analyses, we evaluated the effect of repeated SGS simulations and confirmed that the results sufficiently converged by *n* = 100, stabilizing within a well-defined range of uncertainty (Fig. S2). Examining the spatial distribution of this uncertainty, variance remained higher around the cell periphery, likely due to the lower density of measurement points, making extrapolation challenging (Fig. S3). Nevertheless, the above strong spatial correlation, robust cross-validation performance, and the convergence of SGS estimates justify the use of Pro-FRAP, which applies spatial statistics to estimate unknown diffusion coefficients with statistical rigor.

### 3.2 Spatial analysis of intracellular diffusion coefficients

Four representative raw datasets (Fig. S1), obtained by conventional FRAP measurements at discrete points without spatial statistical processing, were analyzed for subcellular analysis of diffusion coefficients. In all cases, the diffusion coefficients were significantly higher in the cytoplasm compared to the nucleoplasm, with the measurements selectively performed at individually chosen locations during experiments (Fig. 2A, 2B). In contrast, SGS was applied to estimate the diffusion coefficient as a probabilistic variable across the entire cell, using only the data obtained from these same measurement points, resulting in a mapped representation of the mean values (Fig. 2C). For the same representative datasets, regions were classified into the nucleoplasm, the ER, and the rest of the cytoplasm based on the fluorescence intensity of 5-chloromethylfluorescein diacetate, which reflects the extent of probe uptake (Fig. 2D). In all datasets, the diffusion coefficients were highest in the cytoplasm, intermediate in the ER, and lowest in the nucleoplasm (Fig. 2E), a trend consistently observed in individual cells (Fig. S3). These results demonstrate that while traditional FRAP measurements are limited to specific sampling points (Fig. 2A, 2B), Pro-FRAP provides statistically robust diffusion maps that align with trends observed at individual measurement points (Fig. 2C, 2E). This novel approach, therefore, allows for the inference of region-specific characteristics within cells even when the measurement points are limited.

**Figure 2.**
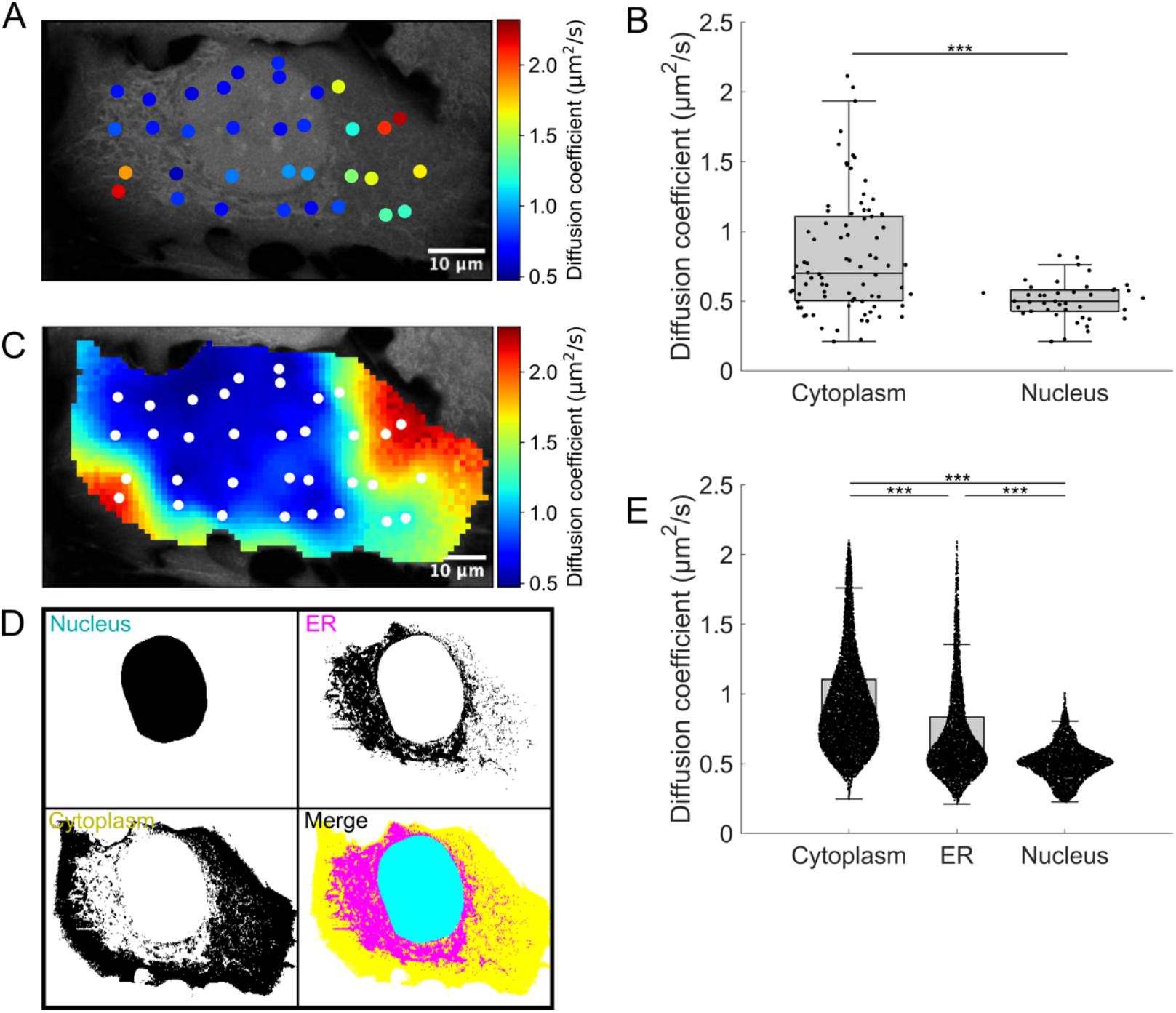
Subcellular analysis of diffusion heterogeneity. (A) Diffusion coefficients measured by FRAP. (B) Boxplots of the diffusion coefficients measured in the cytoplasm (*n* = 82) and nucleus (*n* = 42) from *N* = 4 independent experiments. (C) Diffusion coefficients determined by Pro-FRAP using the same dataset as in (A). White dots represent measurement points. (D) Classification of subcellular regions into the nucleus (cyan), ER (magenta), and the remaining cytoplasmic regions (yellow). (E) Boxplots of the diffusion coefficients for the three classified regions, obtained from the same dataset as in (B), based on *N* = 4 independent experiments.

### 3.3 Numerical simulations evaluate FRAP sampling strategies

The accuracy of intracellular diffusion mapping may be influenced by both the spatial arrangement of measurement points and the underlying diffusion distribution. To systematically investigate these potential factors, numerical simulations were conducted using SGS-based diffusion mapping, incorporating different FRAP sampling patterns and realistic cellular geometries (Figs. 3 and 4). We examined two distinct sampling strategies. In the first case, photobleached positions were determined using a Gaussian function with varying standard deviations *σ* centered within the cell, allowing for a gradual decrease in measurement density from the center to the periphery (Fig. 3A-3C). To characterize these patterns in relation to cellular morphology, an equivalent cell radius *R*_*cell*_ was introduced by approximating the cell as a circle. In the second case, measurement points were arranged in a lattice pattern, ensuring a more uniform spatial coverage across the entire cell (Fig. 3D). The diffusion coefficients were initially modeled as a continuous radial gradient, where values gradually increased from the center to the periphery (Fig. 3A-3D). SGS-estimated values were compared with known ground truths, with MAE and RMSE serving as evaluation metrics. Repeated simulations demonstrated that photobleaching patterns significantly influenced reconstruction accuracy (Fig. 3E). Specifically, prediction errors progressively decreased as *σ* increased, reaching a plateau when *σ*/*R*_*cell*_ exceeded 0.5, a value comparable to half the cell radius. When *σ* was smaller, estimation accuracy declined, likely due to insufficient measurement coverage near the periphery. Similarly, a lattice pattern with a spacing comparable to the *σ*, which corresponds to half the cell radius, improved accuracy (Fig. 3E). Similar trends were observed across different cell morphologies, though elongated cells exhibited greater error variability than rounded ones (Fig. 4A-4C).

**Figure 3.**
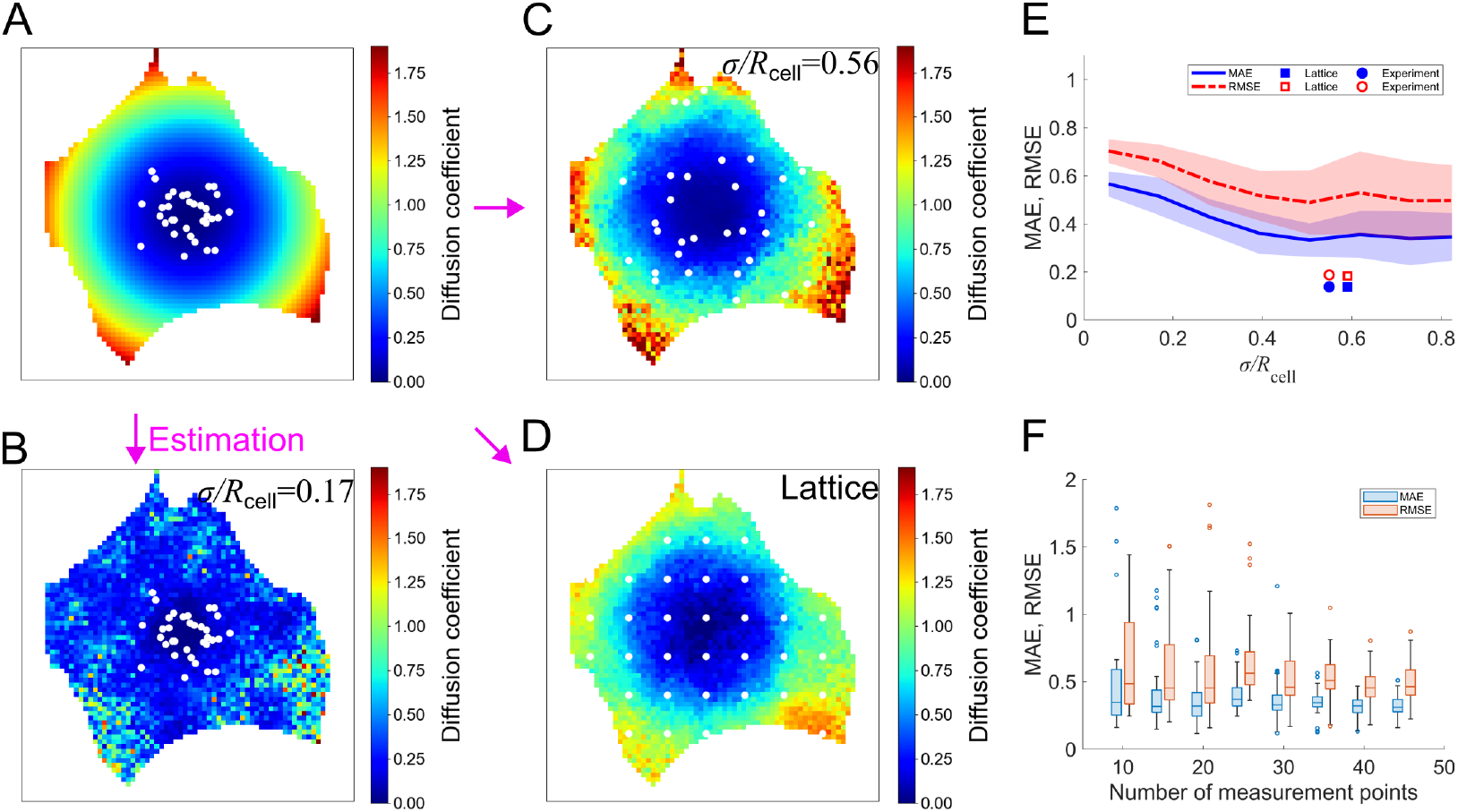
Numerical simulation to evaluate the process of Pro-FRAP. (A) Diffusion coefficients were assigned in a concentric pattern, and measurement points (white dots) were distributed according to a Gaussian function. (B, C) Diffusion coefficients determined by Pro-FRAP at *σ*/*R*_*cell*_ = 0.17 (B) and *σ*/*R*_*cell*_ = 0.56 (C). (D) Diffusion coefficients determined by Pro-FRAP using the same given diffusion distribution as in (A), but with measurement points (white dots) arranged in a lattice pattern. (E) Relationship between errors and *σ*/*R*_*cell*_, where the mean and standard deviation of MAE (blue) and RMSE (red) are shown. The corresponding *σ*/*R*_*cell*_ values for the lattice-based analysis (squares) and those used in actual experimental analysis (circles) are also plotted. (F) Effect of increasing the number of measurement points in SGS analysis on errors (MAE in blue and RMSE in red) when *σ*/*R*_*cell*_ is fixed at 0.56.

**Figure 4.**
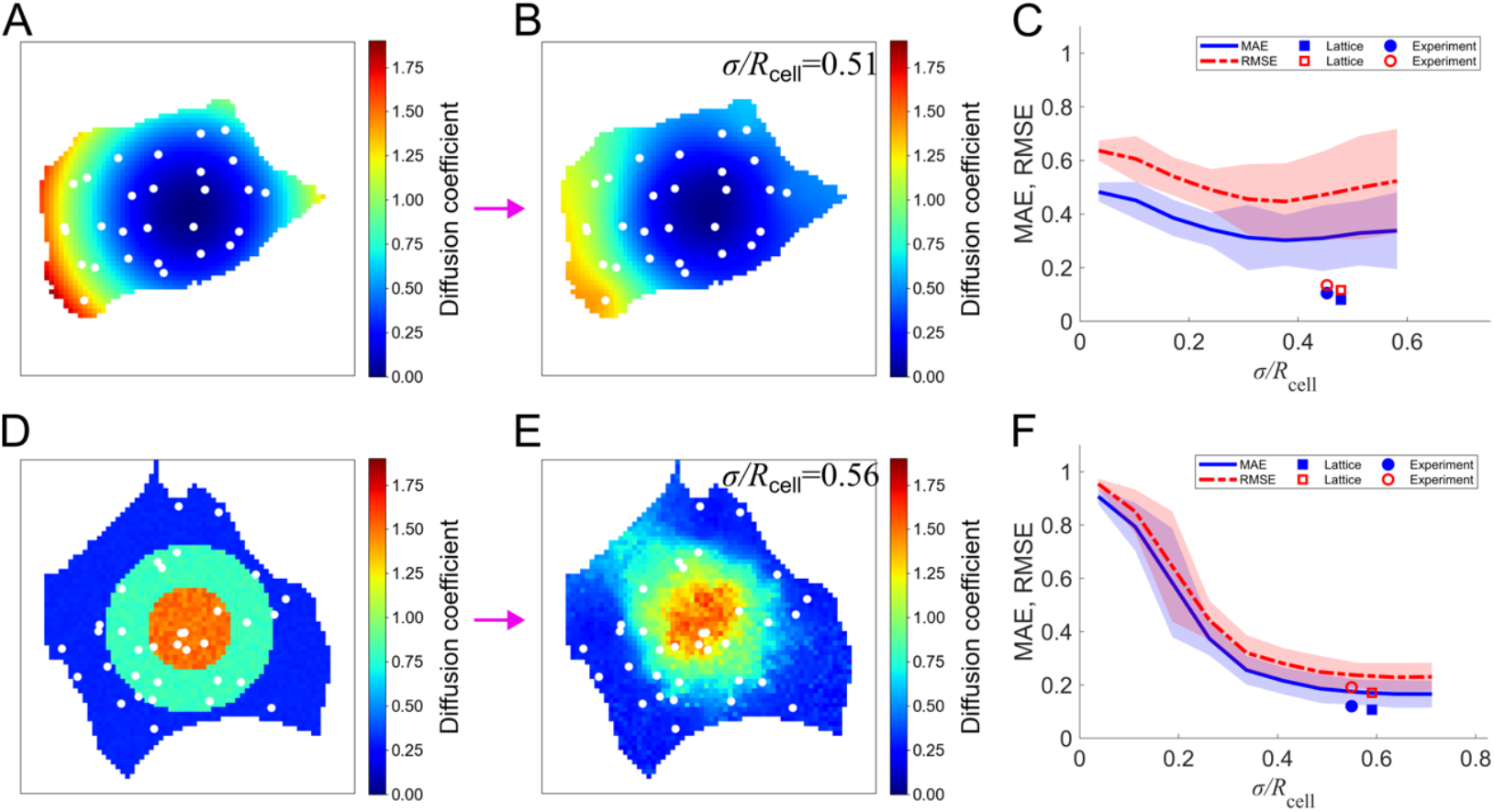
Additional numerical simulation evaluating the performance of Pro-FRAP. (A) Given diffusion coefficient pattern with a continuous concentric gradient and the distribution of measurement points (white). (B) Diffusion coefficients determined by Pro-FRAP with a Gaussian-distributed measurement pattern at *σ*/*R*_*cell*_ = 0.51. (C) Relationship between errors and *σ*/*R*_*cell*_, where the mean and standard deviation of MAE (blue) and RMSE (red) are shown. The corresponding *σ*/*R*_*cell*_ values for lattice-based analysis (squares) and those used in actual experimental analysis (circles) are also plotted. (D) Given diffusion coefficient pattern with stepwise concentric transitions and the distribution of measurement points (white). (E) Diffusion coefficients determined by Pro-FRAP with a Gaussian-distributed measurement pattern at *σ*/*R*_*cell*_ = 0.56. (F) Relationship between errors and *σ*/*R*_*cell*_, where the mean and standard deviation of MAE (blue) and RMSE (red) are shown. The corresponding *σ*/*R*_*cell*_ values for lattice-based analysis (squares) and those used in actual experimental analysis (circles) are also plotted.

To further investigate the effect of measurement density, simulations were conducted under two conditions: ordinary kriging alone and a combination of ordinary kriging with SGS. These analyses were performed using a Gaussian-distributed measurement pattern at *σ*/*R*_*cell*_ = 0.56, with the same cellular morphology and ground truth diffusion coefficient distribution as in Fig. 3A. In ordinary kriging, errors monotonically decreased with increasing measurement points due to its linear interpolation nature, resulting in progressively improved prediction accuracy (Fig. S4). In contrast, although SGS also exhibited error reduction with increasing measurement density, the errors did not converge to zero, while the variability of the estimated values systematically decreased (Fig. 3F). Because SGS sequentially incorporates estimated values as known data, its performance becomes increasingly stable when the initial predictions are accurate. When an optimal FRAP pattern effectively reflects spatial characteristics in the early stages of estimation, subsequent predictions remain reliable, reducing the overall dependence on the number of measurement points. Conversely, suboptimal sampling patterns can introduce spatial inaccuracies that propagate through the sequential estimation process, potentially affecting prediction stability.

In addition, when diffusion coefficients were modeled as a stepwise concentric pattern, where values exhibited discrete transitions rather than a continuous gradient, accuracy stabilized when *σ*/*R*_*cell*_ reached ~0.5 (Fig. 4D-4F), whereas smaller *σ*/*R*_*cell*_ values resulted in larger errors (Fig. S5). These similar trends suggest that the observation remains consistent across different conditions. Accuracy thus consistently stabilized when *σ*/*R*_*cell*_ exceeded ~0.5, corresponding to one-fourth of the cell size. Because a Gaussian distribution extends beyond its standard deviation, measurement points need to cover peripheral regions to maintain accuracy. Based on these findings, we adopted ~30 measurement points per cell in actual experiments, ensuring comprehensive spatial coverage by distributing photobleaching points as evenly as possible.

## Discussion

In this study, we developed Pro-FRAP, a novel framework integrating FRAP with SGS, a powerful spatial statistical method, for estimating molecular diffusion across unmeasured regions. Traditional FRAP techniques, with their localized measurement approach, are inherently limited by excessive photobleaching, which compromises measurement accuracy. Consequently, capturing the full complexity of molecular dynamics within the cellular context becomes challenging. This limitation is particularly significant in analyzing spatial variation given that intracellular macromolecular crowding inherently creates heterogeneity (16).

Spatial statistics have been extensively utilized in various fields including geology, environmental science, and epidemiology to analyze spatially correlated data. These techniques enable the estimation of values in unmeasured regions based on the spatial correlation of measured data points. In our study, we applied the spatial statistical method SGS to analyze intracellular dynamics. As a probabilistic simulation method, SGS enables the quantification of data uncertainty and generates multiple realizations, allowing for a more comprehensive assessment of spatial variability. Our results demonstrated its effectiveness in enabling spatially continuous reconstruction of intracellular diffusion landscapes.

In spatial statistics, unmeasured regions are treated as random variables, enabling statistically robust estimation based on measured data. This approach not only provides a solid statistical foundation but also incorporates spatial autocorrelation, ensuring that predictions align with locally observed data. In contrast, some alternative methods might attempt to determine values in unmeasured regions through deterministic means such as the use of spline functions for interpolation. These methods lack the rigorous statistical justification that spatial statistical techniques offer. By combining the optimal linear unbiased predictions of kriging with the ability of SGS to generate multiple realizations of spatial fields, spatial statistical methods capture the full range of spatial variability while providing statistically grounded predictions.

Other intracellular diffusion measurements such as fluorescence correlation spectroscopy (FCS) also face the same limitation due to their localized measurement nature. While FCS can be strengthened by combining high-speed acquisition to cover larger areas (17), this approach is not feasible for FRAP because extensive photobleaching results in loss of fluorescence signal across the cell. Recent studies have developed the use of small fluorescent particles or genetically encoded multimeric (GEM) nanoparticles to analyze the diffusion coefficient across whole cells (18). However, this method requires customized GEM nanoparticles, and thus it is not applicable to the analysis of specific proteins and other arbitrary molecules. Our approach addresses these challenges by offering a versatile tool that can be adapted to arbitrary fluorescent species without the need for specialized probes.

Although FRAP is traditionally used to measure diffusion, it can also provide insights into other dynamic processes such as chemical binding and dissociation, namely turnover, as well as macroscopic deformations and advection (19, 20). Although we focused herein only on the diffusion of a low-molecular-weight fluorescent compound to evaluate the usefulness of our approach, the combination of FRAP with spatial statistical techniques may be extended to analyze such other physicochemical properties, which will be the subject of future research. In this regard, while the recently developed method using GEM nanoparticles is limited to diffusion measurement, our approach potentially covers more diverse applications, providing a more comprehensive understanding of the complex molecular behavior in cells.

The complexity of the intracellular environment necessitates advanced statistical methods for accurate analysis. Our separate analyses revealed that while kriging alone achieves an average prediction accuracy of ~75%, SGS displayed superior accuracy, achieving ~85% (Fig. S1). Kriging optimally utilizes spatial correlations for predictions with well-defined statistical confidence, but its reliance on nearby observations can limit its ability to capture global variability. Indeed, increasing the number of measurement points consistently improved accuracy (Fig. S4), but this approach is impractical due to fluorescence bleaching constraints. In contrast, SGS effectively captures spatial variability with lower error fluctuation, though it requires a substantial computational load. Despite this, the advantages of SGS allowed us to achieve high-resolution, statistically reliable mapping of molecular diffusion within cells.

Recent advances in omics technologies and mass spectrometry, including single-cell RNA sequencing, mass spectrometry imaging, and spatial transcriptomics, have revolutionized the spatial mapping of intracellular components at a single-cell level, providing unprecedented insights into cellular spatial organization (21–24). Combining these cutting-edge technologies with the Pro-FRAP approach could offer new perspectives on how intracellular components and their physicochemical dynamics are functionally interconnected. Thus, the proposed method has the potential to deepen our understanding of cellular functions and their spatial regulation.

In conclusion, Pro-FRAP, which combines FRAP with spatial statistical analysis, provides a robust framework for visualizing molecular diffusion at the whole-cell level. This approach addresses the inherent limitations of traditional FRAP techniques, particularly those related to localized measurements and photobleaching. By integrating the advanced spatial statistics method SGS, we achieved statistically grounded mapping of molecular diffusion. This versatile framework is not inherently limited to diffusion mapping alone but can be extended to analyze other dynamic processes such as turnover, macroscopic deformations, and advection, ultimately providing deeper insights into the complex physicochemical behavior of intracellular molecules.

## Author contributions

YO performed the experiments with feedback from TS. YO analyzed the data with feedback from TS and DM. ON conducted the SGS analysis. YO and ON prepared the figures with feedback from TS and SD. DM provided support for analysis and resources. SD led the design of the research and supervised its execution. TS and SD wrote the draft. All authors reviewed and approved the final manuscript.

## Acknowledgments

This study was partly supported by JSPS KAKENHI grants (21H03796, 22J00060, 23H04929).

## Supporting materials

**Figure S1.**
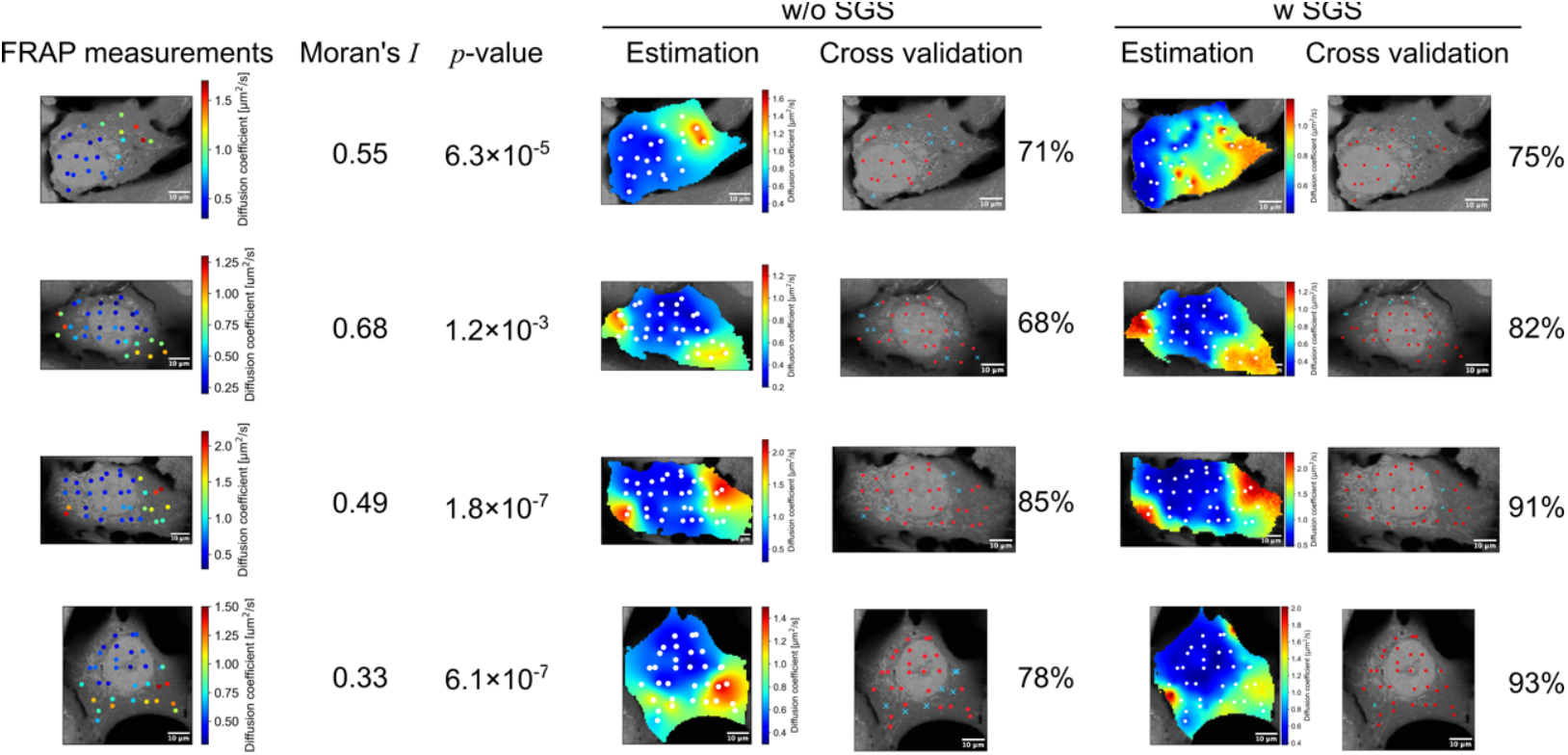
Validation of Pro-FRAP using four representative datasets. The left column shows the diffusion coefficients measured by FRAP, along with the corresponding Moran’s *I* and *p*-values. The middle column (w/o SGS) and the right column (w SGS) display the diffusion coefficient distributions determined using kriging alone (middle) and Pro-FRAP with SGS (right), respectively. Cross-validation results are also shown, where correctly predicted points are marked in red, and incorrectly predicted points defined as those deviating by more than one standard deviation from the normal distribution are indicated by blue crosses.

**Figure S2.**
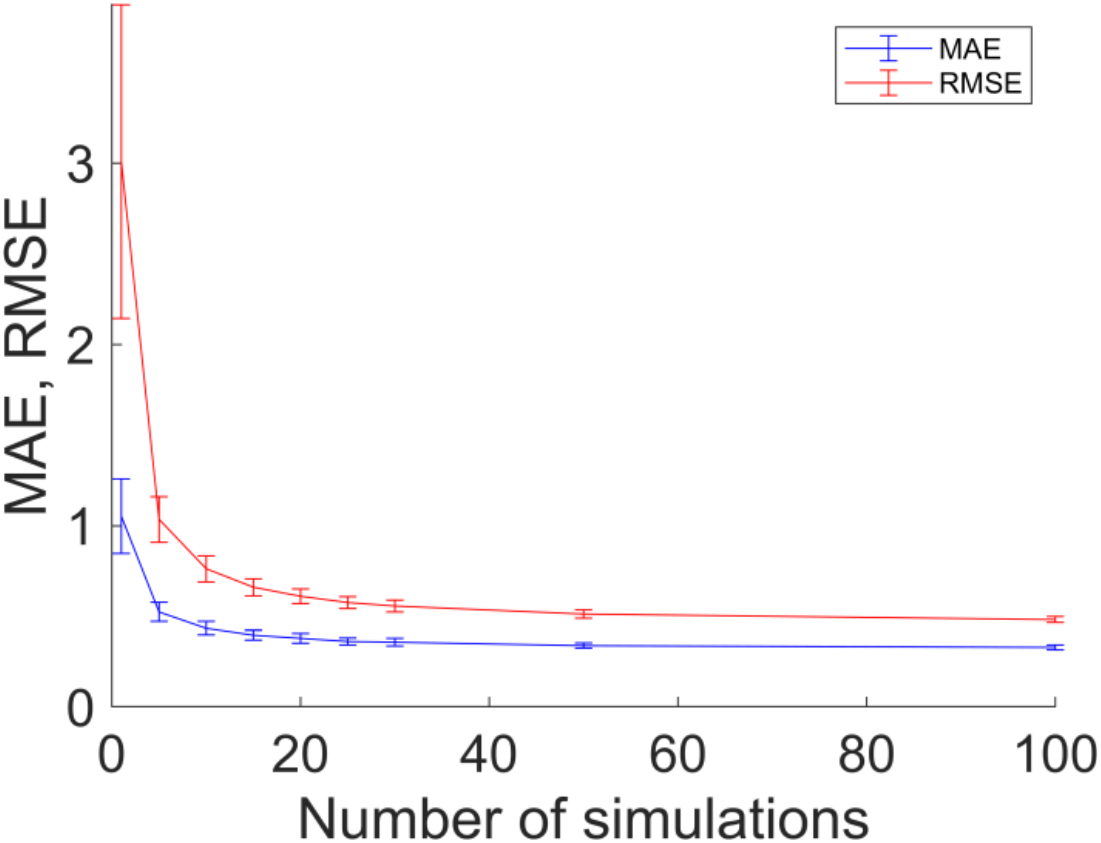
Effect of the number of SGS iterations on error convergence. Both MAE and RMSE sufficiently converge by *n* = 100.

**Figure S3.**
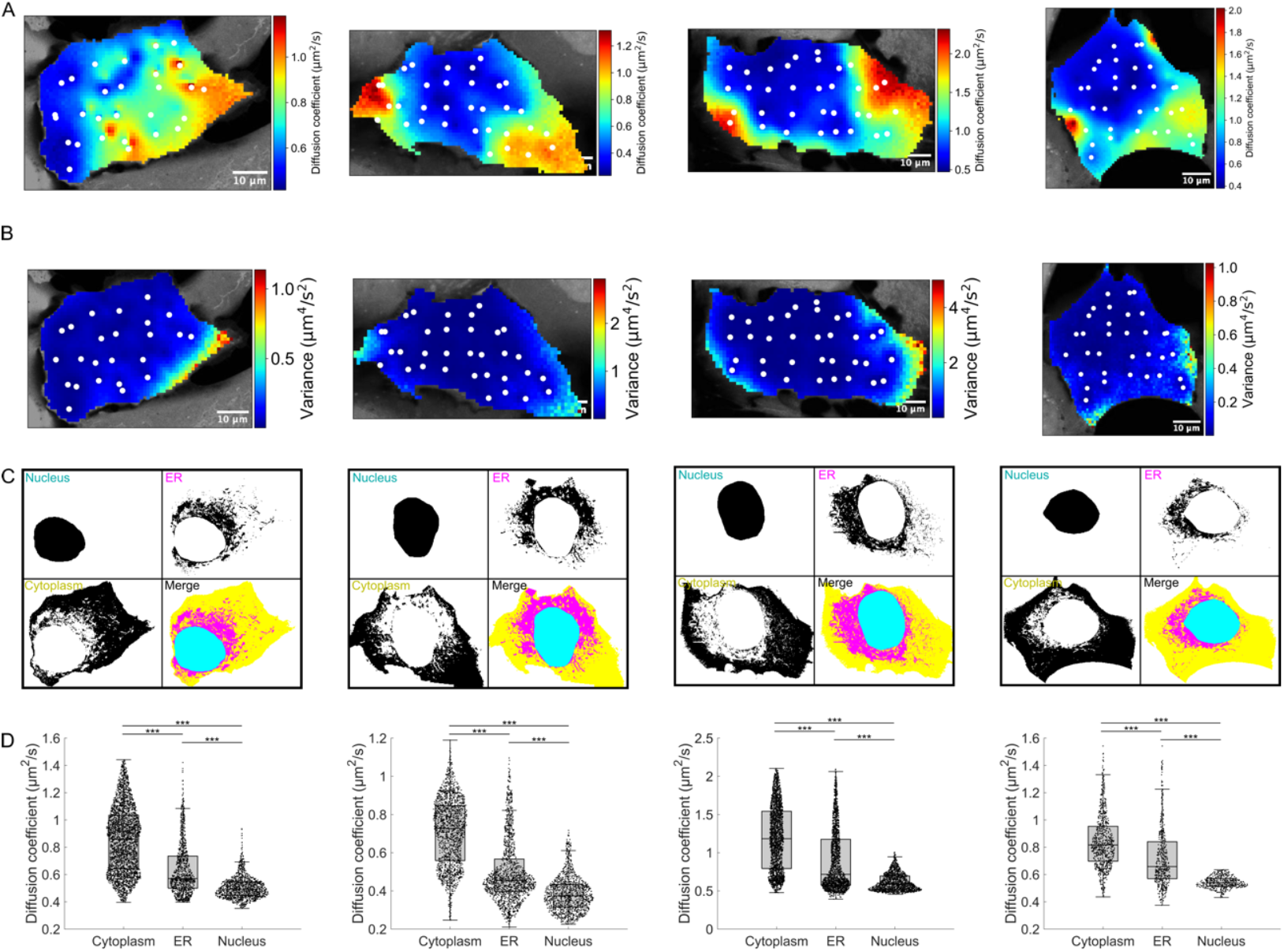
Subcellular analysis of diffusion coefficients for four representative datasets. (A, B) Mean diffusion coefficients (A) and variance (B). (C) Segmentation of subcellular regions into the nucleus, ER, and remaining cytoplasm. (D) Diffusion coefficients for each classified region.

**Figure S4.**
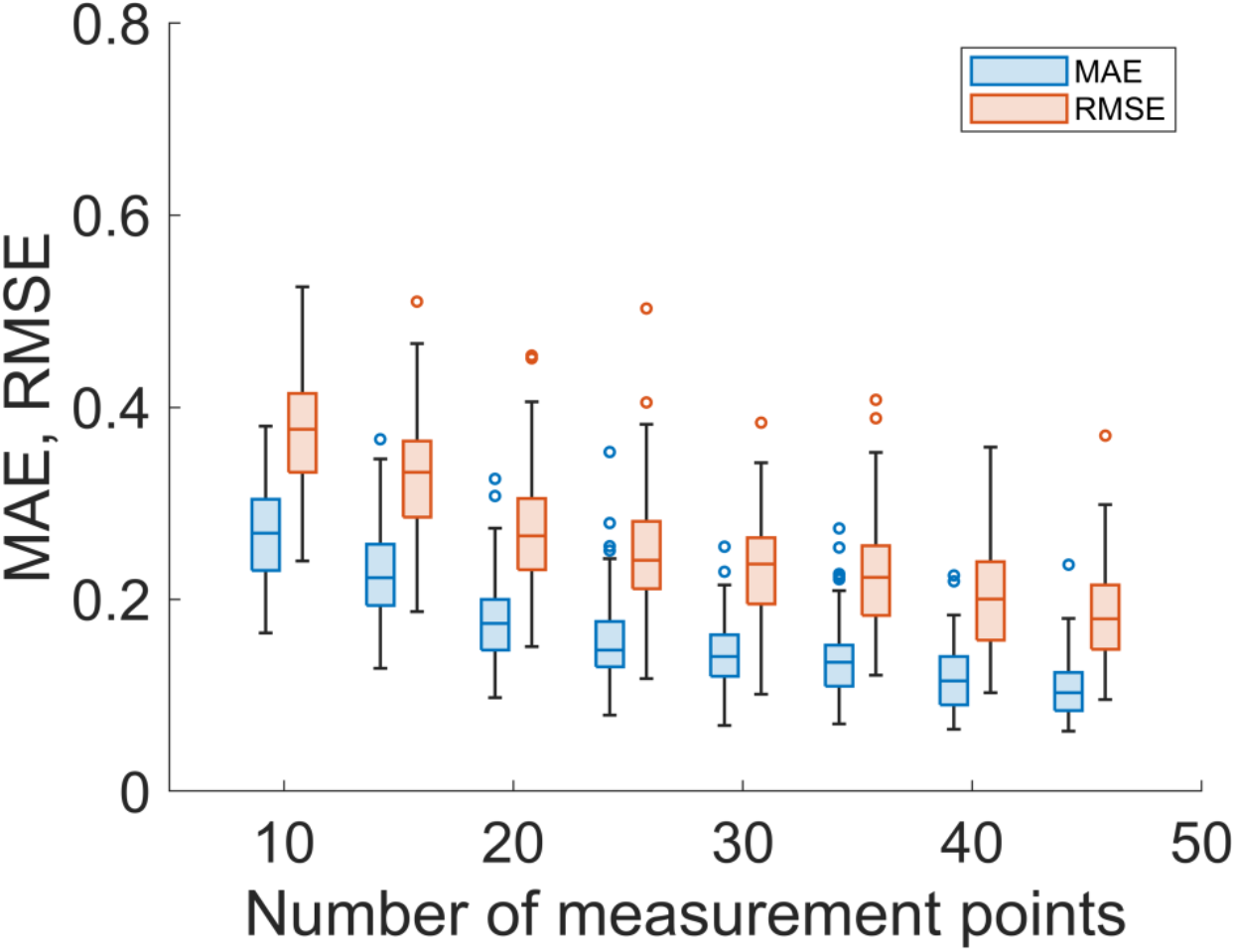
Evaluation of ordinary kriging. Effect of increasing the number of measurement points in ordinary kriging alone on errors (MAE in blue and RMSE in red) when *σ*/*R*_*cell*_ is fixed at 0.56.

**Figure S5.**
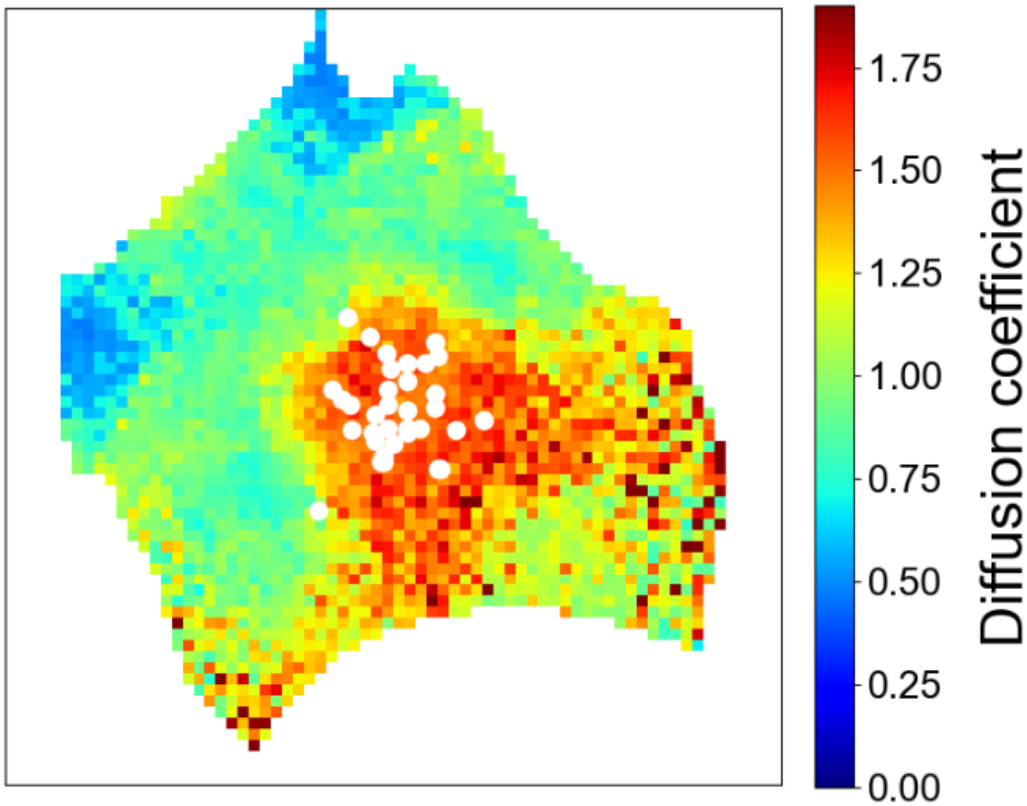
Additional analysis of Fig. 4D. Diffusion coefficients determined by Pro-FRAP with a Gaussian-distributed pattern of *σ*/*R*_*cell*_ = 0.17, corresponding to Fig. 4D.

## Supporting note 1

Ordinary kriging assumes a steady state in the spatial domain, and the mean of a physical variable *m* is constant but unknown. The unbiasedness condition for predictions is expressed as

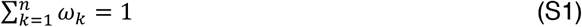

Here, *ω*_*k*_ represents the weights assigned to the observed data positions *u*_*k*_. The prediction error variance of a variable *Z* relative to the true value is expressed as

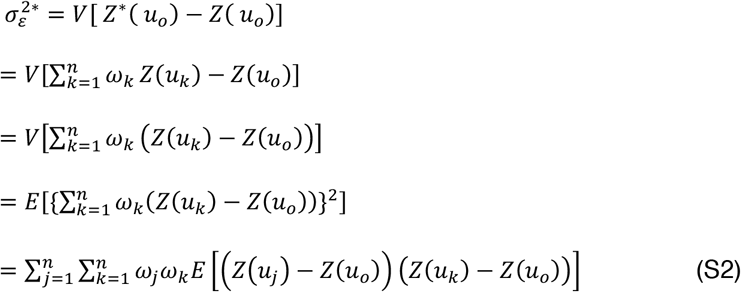

where *V* and *E* represent the variance and expected value, respectively; *u*_*o*_ and *u*_*k*_ denote the positions of an unobserved point and of an arbitrary point, respectively; and an asterisk indicates estimated variables. The semi-variogram is defined as half of the squared difference between values at any given distance *h*:

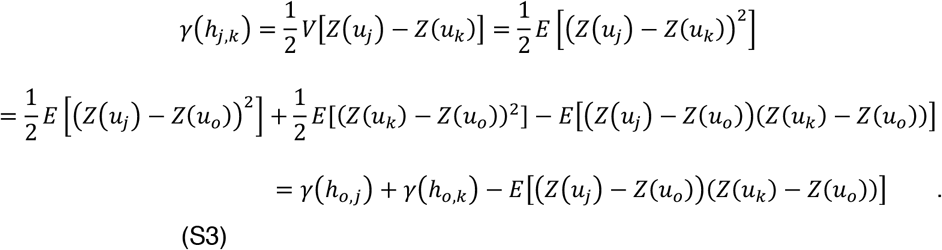

Substituting Eq. (S3) into Eq. (S2) and eliminating *E*,

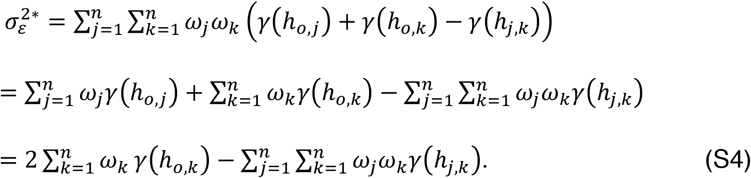

We consider a Lagrange multiplier problem to minimize the semi-variogram under the constraint given by Eq. (S1),

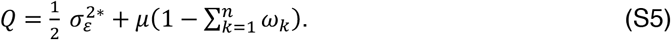

The variables *ω*_*k*_ and *µ* are determined by minimizing the objective function *Q*, and thus

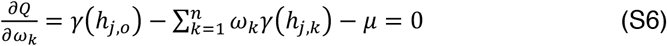

and

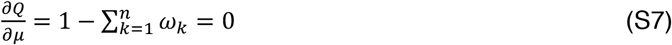

The combination of Eqs. (S6) and Eq. (S7) forms the kriging equations (i.e., Eq. (3) and (4) in the main text), which are a system of *n* + 1 linear equations with unknowns *ω*_*k*_ (*k* = 1, 2, …, *n*) and *µ*.

